# Expression graph network framework for biomarker discovery

**DOI:** 10.1101/2025.04.28.651033

**Authors:** Yang Liu, Jason Huse, Kasthuri Kannan

## Abstract

Biomarker discovery for complex diseases like cancer hinges on uncovering molecular signatures that capture intricate, interconnected relationships within biological data, a challenge that traditional statistical and machine learning methods often fail to meet due to the complexity of high-dimensional gene expression profiles. To overcome this, we introduce the Expression Graph Network Framework (EGNF), a cutting-edge graph-based approach that integrates graph neural networks (GNNs) with network-based feature engineering to enhance predictive biomarker identification. EGNF constructs biologically informed networks by combining gene expression data and clinical attributes within a graph database, utilizing hierarchical clustering to generate dynamic, patient-specific representations of molecular interactions. Leveraging graph learning techniques, including Graph Convolutional Networks (GCNs) and Graph Attention Networks (GATs), our framework identifies statistically significant and biologically relevant gene modules for classification. Validated across three independent datasets consisting of contrasting tumor types and clinical scenarios, EGNF consistently outperforms traditional machine learning models, achieving superior classification accuracy and interpretability. Notably, it delivers perfect separation between normal and tumor samples while excelling in nuanced tasks such as classifying disease progression and treatment outcome. This scalable, interpretable, and robust framework provides a powerful tool for biomarker discovery, with wide-ranging applications in precision medicine and the elucidation of disease mechanisms across diverse clinical contexts.

## 1 Introduction

Classification problems constitute the cornerstone of machine learning, with the paramount objective of categorizing observations into discrete classes based on their attributes. This process is especially critical in healthcare, where precise classification directly influences diagnosis, treatment strategies, and patient outcomes. Machine learning has revolutionized conventional analytical methods, offering innovative approaches to process and interpret complex datasets. The application of these techniques to biomarker discovery has emerged as a particularly promising domain, providing researchers with powerful tools to identify molecular indicators that can predict disease states, progression, and treatment response.

Graph-based learning approaches have garnered significant attention in biomedical research due to their distinctive ability to model intricate relationships between biological entities. Unlike traditional machine learning methods that treat samples as independent observations, graph-based approaches leverage the inherent interconnectedness of biological data, capturing relationships that might otherwise remain obscured. This capability is particularly valuable in biomarker discovery, where understanding the interactions between molecules can provide more profound insights into disease mechanisms than analyzing individual features in isolation.

Cluster-based feature engineering represents a powerful technique for improving machine learning models by grouping similar data points based on shared attributes, enabling extraction of meaningful patterns. One widely used method is the k-means clustering algorithm, which clusters data points based on proximity in feature space (MacQueen, 1967). K-means clustering has been employed in various feature engineering applications to enhance model performance by identifying relationships between neighboring data points. By analyzing inter-relationships within these clusters, models can make more informed predictions, as interactions between neighboring data points often reveal underlying data structures (Mostafa & Amano, 2019). Unlike dimensionality reduction techniques, cluster-based methods like k-means clustering focus on enhancing model performance by capitalizing on these inter-relationships rather than reducing feature count. Hierarchical clustering extends this concept by creating nested clusters at different levels of similarity, which aligns conceptually with the multi-level nature of biological interactions and provides a natural framework for discovering biomarker relationships across different scales of biological organization.

Among various classification techniques, conventional methods like logistic regression remain notable for their simplicity and interpretability in biomarker studies. Despite their widespread utilization, these methods frequently fail to capture the complex, non-linear relationships present in biological systems, constraining their utility for comprehensive biomarker discovery. While Support Vector Machines (SVMs), random forest models, and elastic net regression offer more robust approaches to biomarker discovery, especially in high-dimensional settings, these methods primarily operate on tabular data and do not inherently account for the network structure of biological systems. Random forest models, though capable of capturing non-linear interactions through their ensemble of decision trees, still treat features independently and cannot directly incorporate the relational information encoded in biological networks.

GNNs have emerged as a powerful class of models designed to advance biomarker discovery by leveraging graph-structured data prevalent in biological applications. GNNs capture complex relationships between biological entities represented as nodes, along with their interactions defined by edges. Among the most prominent GNN architectures are GCNs and GATs. GCNs extend convolutional neural networks to graph data by leveraging the adjacency structure, enabling efficient information propagation among connected features (Kipf & Welling, 2017). GATs enhance this capability by introducing attention mechanisms that allow the model to weight the importance of different node neighbors dynamically (Veličković et al., 2018), while more recent architectures further improve performance through positional and structural encodings that better capture graph topology (Brody et al., 2021).

The integration of multi-omics data through graph-based approaches has significantly enhanced identification of clinically relevant biomarkers. Previous studies have demonstrated superior performance in patient classification and biomarker identification compared to conventional methods. For instance, Wang et al. (2021) introduced MOGONET, a framework integrating multi-omics data using graph convolutional networks. Similarly, Ramirez et al. (2020) applied GCNs to cancer classification, showcasing the potential of graph-based approaches in distinguishing between cancer types based on gene expression data. These studies highlight the power of graph-based methods in capturing the complex interplay between different biological layers.

Existing graph-based approaches for biomarker discovery typically rely on established biological networks, such as protein-protein interaction networks or co-expression networks. The Weighted Gene Co-expression Network Analysis (WGCNA) developed by Langfelder and Horvath (2008) has been widely used to identify modules of co-expressed genes that may serve as potential biomarkers. Building upon this foundation, recent approaches have proposed multi-level attention graph neural networks based on co-expression gene modules for disease diagnosis and prognosis, demonstrating improved predictive performance and interpretability.

Despite these advances, a critical limitation of existing approaches is that they are not specifically tailored for tissue sample classification in biomarker discovery. Most methods rely on pre-defined biological networks, which may not accurately reflect the specific relationships relevant to the disease or condition under investigation. Additionally, these approaches often struggle to handle datasets with varying sample sizes, limiting their applicability across different clinical contexts. Therefore, we introduce EGNF, a novel methodology integrating network generation with GCNs and GATs for gene expression-based classification.

This methodology advances machine learning in biomedical applications through two primary contributions. First, EGNF leverages deep learning to enhance the extraction of complex patterns and relationships from gene expression data, significantly improving classification accuracy. While deep learning has been applied for these purposes, our approach uniquely employs hierarchical clustering to identify meaningful biological relationships, providing a natural bridge between conventional cluster-based feature engineering and advanced graph-based learning methods. The generated networks capture intricate relationships between samples and features, adaptively configuring to different sample sizes while preserving biological relevance.

Second, we establish a biologically meaningful network-based feature selection method tailored to gene expression data. By combining network analysis with conventional statistical techniques, our method identifies gene modules that are both statistically significant and biologically relevant, offering deeper insights into disease progression mechanisms. This approach reduces data complexity while maintaining predictive power and ultimately improves the interpretability of machine learning models.

Together, these contributions offer several advantages over existing methods: (1) they enable more accurate patient/sample stratification by leveraging complex patterns encoded in generated graphs; (2) they provide insights into biological mechanisms underlying disease states by highlighting important connections between biomarkers; (3) they facilitate integration of multi-modal data, capturing relationships spanning different biological domains; and (4) they demonstrate robust performance across different datasets and disease types, suggesting broad applicability in precision medicine.

## 2 Methods

### 2.1 Overview of EGNF

Our methodological framework consists of several sequential analytical stages (Figure 1). Initially, we performed differential expression analysis on 80% of the data using DESeq2 to identify differentially expressed genes (DEGs) (Love et al., 2014). Using this training data, we constructed a graph network by selecting extreme sample clusters with high or low median expression for each group (unpaired method), or group ratio values (paired method) from one-dimensional hierarchical clustering as nodes and establishing connections between sample clusters of different genes through shared samples. We then conducted graph-based feature selection considering three criteria: node degrees, gene frequency within communities, and inclusion in known biological pathways. The selected features were then used to generate sample clusters via one-dimensional hierarchical clustering, which served as nodes for building the prediction network. In the final stage, we employed GNNs for sample-specific graph-based predictions, representing each sample by its corresponding subgraph structure.

**Figure 1:**
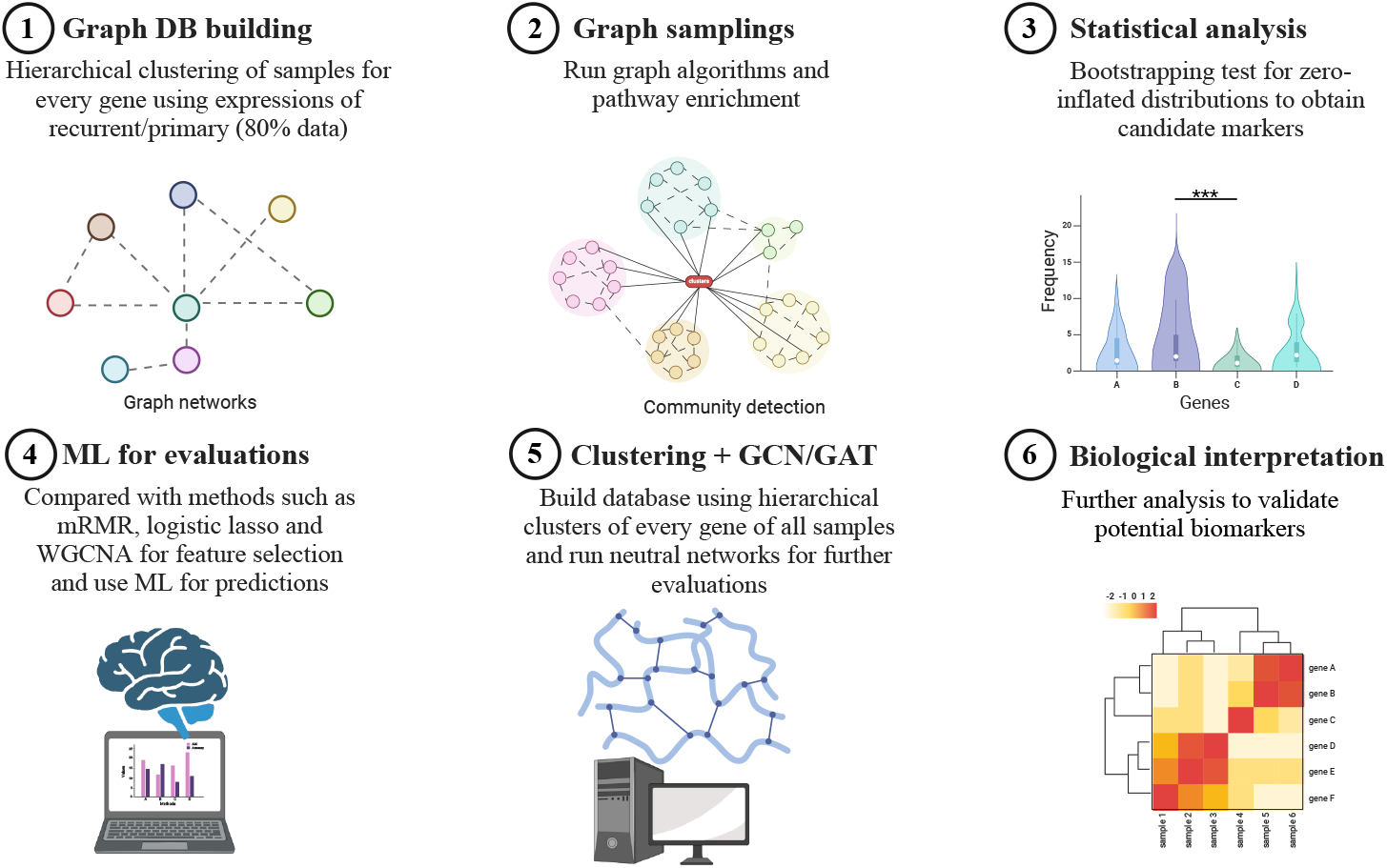
Framework for network-based feature selection

This study utilized open-source libraries for biomarker discovery, including PyTorch Geometric for GNN model development and network analysis tools, such as Neo4j and GDS. All algorithm development and validation phases were executed within this computational framework to ensure reproducibility and scalability.

### 2.2 Datasets

This study used three paired gene expression datasets to assess model performance across clinical contexts. The glioma dataset comprised 295 primary and 275 corresponding recurrent tumors from IDH-WT patients, sourced from the Glioma Longitudinal AnalySiS Consortium (GLASS) and MD Anderson Cancer Center’s Glioblastoma Moon Shot project, enabling analysis of molecular changes during disease progression. The breast cancer dataset included 111 normal tissue specimens paired with 113 matched tumor samples from The Cancer Genome Atlas Program (TCGA), allowing examination of differences between healthy and malignant tissues. The third dataset contained 69 matched pre- and post-treatment samples from HER2-negative breast cancer patients who underwent neoadjuvant chemotherapy with Bevacizumab (GSE87455), obtained from the GEO repository, facilitating analysis of treatment-induced molecular alterations. All datasets exclusively included patients with complete paired gene expression profiles, with unpaired samples excluded from analysis.

### 2.3 Network Generation

We employed a bottom-up one-dimensional hierarchical clustering approach for each gene to identify sample clusters, using Euclidean distance as the similarity metric (Figure 2). Clusters were merged according to the median linkage criteria, which calculates inter-cluster distances based on the median of pairwise distances between cluster centroids. The resulting hierarchical trees were imported into a Neo4j graph database to construct gene-gene interaction networks. Each node represents a gene with associated properties, including expression level, gene identifier, sample count, and median expression value, in this network representation. Edges between nodes represent shared samples between gene clusters, with the number of common samples serving as an adjustable parameter (default threshold: 1). Properties including the common sample count and the respective levels of the connected nodes characterized each edge.

**Figure 2:**
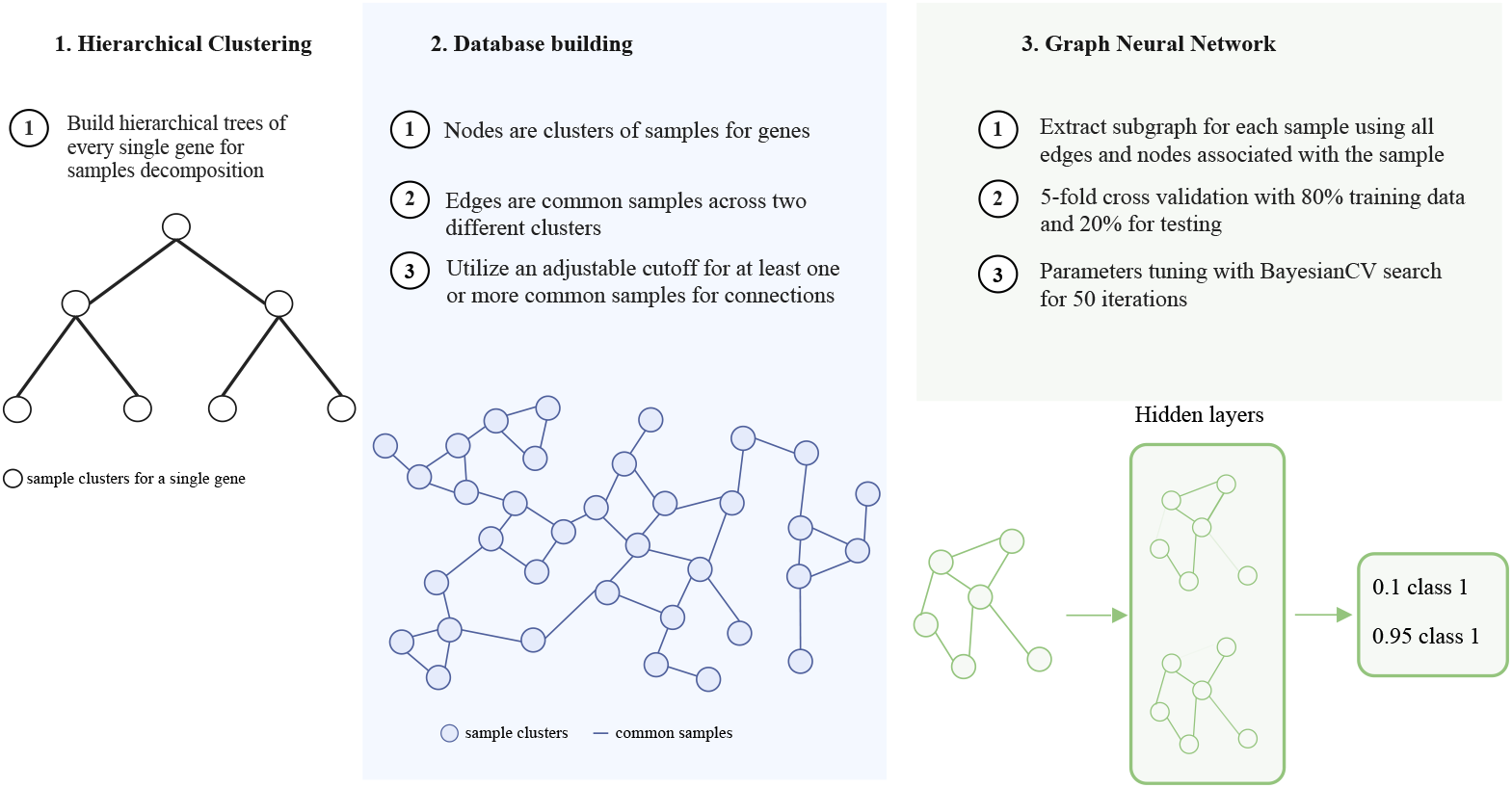
Network generation process

The Euclidean distance between two samples *x*_*i*_ and *x*_*j*_ is given by:

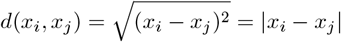

### 2.4 Graph-Based Feature Selection for Identifying Biomarkers

We developed two network-based feature selection methods for identifying significant biomarkers in paired and unpaired datasets. Both approaches begin with dimension reduction using DESeq2, followed by dataset-specific normalization. For paired datasets, we applied log_2_(*x* + 1) normalization followed by rescaling values to a range of 1 to 2 for each sample class, then used class2/class1 ratios to perform one-dimensional hierarchical clustering. For unpaired datasets, we employed log_2_(*x* + 1) normalization followed by z-normalization for each sample class and conducted hierarchical clustering separately. The top 10% of sample/patient clusters with either high or low expression values for each gene form network nodes, with shared patients defining edges to focus on gene-gene interactions.

To reduce the impact of indirect, complex relationships among the same gene, we randomly selected a single sample cluster for each gene per graph sampling iteration and repeated this process 10,000 times. Degree centrality algorithms were applied to enumerate degrees for each gene. Modularity optimization algorithms were utilized to delineate gene communities. For each community, we quantified gene frequency and scrutinized whether genes within communities are incorporated in enriched pathways. Bootstrapping analyses further refined marker selection by comparing each marker’s mean expression against the mean expression of all alternative markers, addressing the zero-inflated distribution of degree, frequency, and pathway scores.

To evaluate the impact of pathway enrichment on predictive performance, we constructed reference marker catalogues devoid of pathway enrichment. A comprehensive scoring system was devised to prioritize markers by ranking adjusted p-values across various filtration criteria and summing ranks for definitive selection.

### 2.5 Other Feature Selection Methods

WGCNA, ANOVA, and mRMR are three complementary feature selection methods used to identify informative genes from high-dimensional gene expression data. WGCNA constructs gene co-expression networks by clustering genes based on their correlation patterns, identifying modules of highly correlated genes, and associating them with phenotypic traits to uncover biologically relevant relationships. ANOVA, a statistical approach, determines whether gene expression levels significantly differ across multiple groups by comparing within-group and between-group variability. Lastly, mRMR selects features that are both highly relevant to the target variable and minimally redundant with each other, ensuring an informative yet non-redundant subset of genes for downstream analysis. These methods collectively enhance the robustness of feature selection by integrating network-based, statistical, and information-theoretic approaches.

### 2.6 Graph Convolutional Network (GCN) Model

GCNs extend the concept of convolutional operations to graph-structured data. A GCN model operates by aggregating feature information from neighboring nodes in a graph to learn node representations. Given a graph *G* = (*V, E*) where *V* represents the set of nodes and *E* the set of edges, let *A* be the adjacency matrix. The propagation rule of a GCN layer can be expressed as:

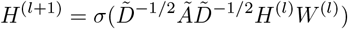

where *Ã* = *A* + *I*_*N*_ is the adjacency matrix with added self-connections,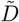 is the diagonal degree matrix, *H*^(*l*)^ is the node feature matrix at layer *l, W* ^(*l*)^ is the learnable weight matrix at layer *l*, and *σ* ( *·* ) is a non-linear activation function such as ReLU.

### 2.7 Graph Attention Network (GAT) Model

GATs introduce attention mechanisms into GNNs, allowing the model to assign different importance weights to different neighbors. The attention coefficient between node *i* and neighbor *j* is computed as:

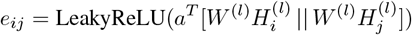

where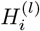and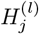are the input feature vectors of nodes *i* and *j* for layer *l, W* ^(*l*)^ is a learnable weight matrix, *a* is a learnable attention vector, denotes concatenation, and LeakyReLU is an activation function. The attention coefficients are normalized using the softmax function across all neighbors *j* of node *i*:

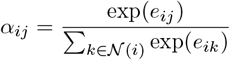

The node feature update for node *i* is then computed as a weighted sum of its neighbors’ features:

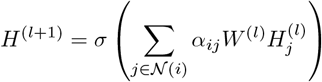

Where *σ* here is a non-linear activation function, and *α*_*ij*_ are the attention coefficients that weight the contribution of each neighbor’s features. GCNs aggregate neighborhood information uniformly using normalized adjacency matrices, while GATs utilize an attention mechanism to focus on the most relevant neighbors, dynamically weighting each neighbor’s contribution to a node’s feature representation.

### 2.8 Graph Attention Network v2 (GATv2) Model

GATv2 improves upon the original GAT by introducing dynamic attention mechanisms that better capture node relationships (Brody et al., 2021). Unlike GAT, where the attention coefficients are computed in a static manner, GATv2 allows for a more expressive and adaptive attention function by making the attention mechanism order-invariant. This enables the model to assign attention to weights dynamically based on node features rather than relying on predefined structures.

The attention mechanism in GATv2 is defined similarly to GAT but introduces a key modification. For a node, the attention coefficient between node and one of its neighbors is computed as:

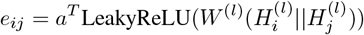

Beyond this modification, the normalization of attention coefficients and the node feature update follow the same formulation as in GAT. By leveraging this improved attention mechanism, GATv2 provides better adaptability and expressiveness in learning from graph-structured data.

### 2.9 Traditional Machine Learning Models

Traditional machine learning approaches have shown considerable success in various classification tasks. Logistic regression, a fundamental statistical model, provides interpretable results by modeling probability through a logistic function (Hosmer et al., 2013). The elastic net extends conventional regression by incorporating both L1 and L2 regularization, effectively handling multicollinearity and performing feature selection simultaneously (Zou & Hastie, 2005). Random Forest, an ensemble learning method, combines multiple decision trees to reduce overfitting and improve generalization by leveraging bagging and random feature selection (Breiman, 2001). SVM excels in finding optimal hyperplanes to separate classes in high-dimensional spaces through kernel transformations (Cortes & Vapnik, 1995). MLP, a type of artificial neural network, can capture complex non-linear relationships by utilizing multiple layers of interconnected neurons with activation functions (Rumelhart et al., 1986). These models have been widely applied across various domains, demonstrating their versatility and effectiveness in handling different types of data and classification problems.

### 2.10 Parameter Tuning Process

Model hyperparameters were optimized using 5-fold cross-validation on the training dataset. For conventional machine learning models (logistic regression, elastic net, random forest, SVM, and MLP), we employed random search to efficiently explore parameter spaces (Bergstra & Bengio, 2012), optimizing regularization parameters for elastic net, tree parameters for random forest, kernel parameters for SVM, and network architecture for MLP. For GNNs, we utilized Bayesian optimization to navigate their more complex parameter spaces, including network depth, learning rates, and attention mechanisms (Snoek et al., 2012). Conventional machine learning analyses were implemented using the caret package (Kuhn, 2008) in R with parallel processing, while GNN computations were performed on an NVIDIA GPU with 45 GB VRAM in an HPC environment. All models were evaluated through ten iterations of training and testing, with final performance assessed using average accuracy and AUC scores.

## 3 Results

### 3.1 GNN-based classification improves performance

The machine learning models, trained on gene expression profiles selected through WGCNA, mRMR, and ANOVA, displayed varied performance across different classification tasks (Figure 3). They showed strong results in binary classification of normal versus tumor samples in breast cancer, achieving high accuracy and AUC. However, their performance decreased in more complex tasks like distinguishing primary from recurrent tumors and pre-from post-treatment samples.

**Figure 3:**
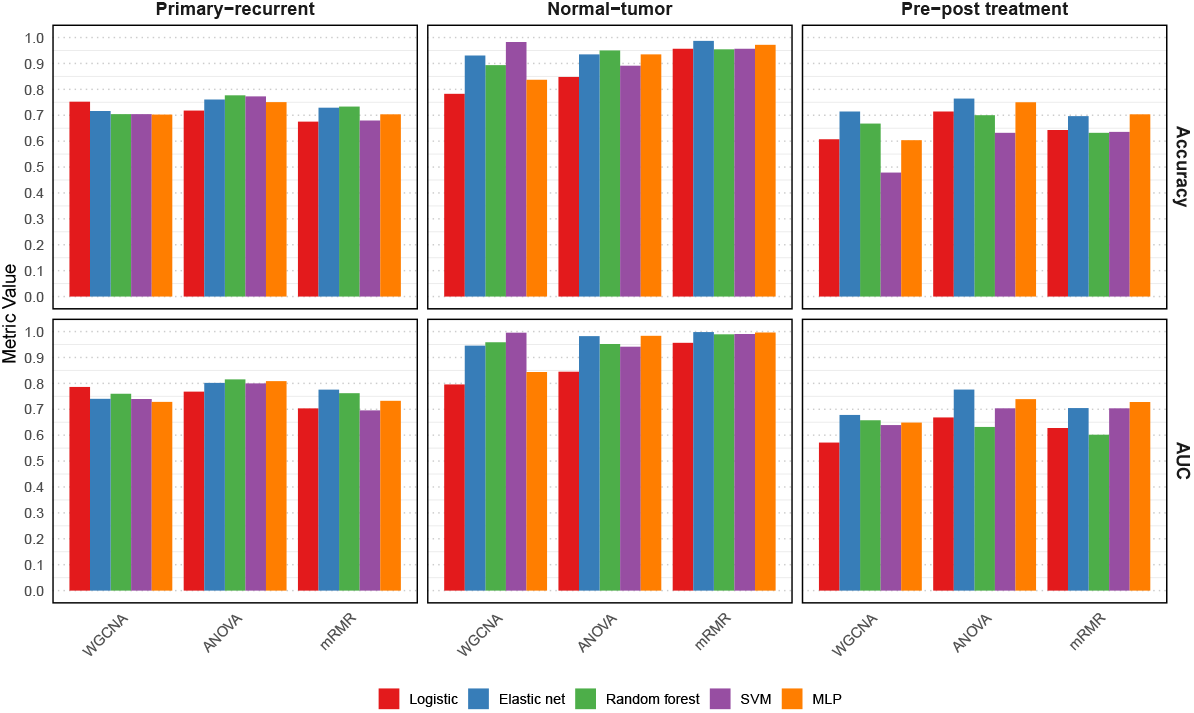
Performance of machine learning models for different features

We implemented our methodology incorporating network generation and GNNs and compared its performance against traditional machine learning models (Figure 3) using accuracy and AUC as baselines across 10 iterations. Table 1 presents the relative improvements achieved by our GNN-based classification for each feature set. Our method enhanced accuracy by 13.8%, -2.4%, and 2.0% across the three datasets. Similarly, it improved AUC by 17.1%, -0.7%, and 1.3%, respectively. The variability in magnitude improvement is likely due to differences in sample sizes and the complexity of the classification task. GNNs faced challenges in achieving superior performance in data-limited contexts.

**Table 1:** Performance comparison across datasets and feature selection methods.

| Dataset | Features | Accuracy (SD) |  |  |  | AUC (SD) |  |  |  |
| --- | --- | --- | --- | --- | --- | --- | --- | --- | --- |
|  |  | GCN-gene | GCN | GAT | GATv2 | GCN-gene | GCN | GAT | GATv2 |
| Primary-recurrent | WGCNA | 0.717 (0.017) | 0.748 (0.058) | 0.698 (0.063) | 0.740 (0.008) | 0.780 (0.013) | 0.782 (0.085) | 0.731 (0.088) | 0.771 (0.011) |
|  | ANOVA | 0.838 (0.006) | 0.847 (0.034) | 0.855 (0.017) | 0.850 (0.020) | 0.901 (0.005) | 0.897 (0.032) | 0.901 (0.019) | 0.900 (0.024) |
|  | mRMR | **0.884 (0.010)** | 0.820 (0.056) | 0.866 (0.021) | 0.862 (0.020) | **0.954 (0.004)** | 0.892 (0.055) | 0.932 (0.018) | 0.924 (0.009) |
| Normal-tumor | WGCNA | 0.813 (0.027) | **0.963 (0.023)** | 0.917 (0.030) | 0.922 (0.026) | 0.806 (0.014) | **0.991 (0.008)** | 0.961 (0.020) | 0.967 (0.017) |
|  | ANOVA | 0.583 (0.073) | 0.943 (0.046) | 0.865 (0.132) | 0.913 (0.023) | 0.399 (0.191) | 0.978 (0.032) | 0.907 (0.144) | 0.950 (0.015) |
|  | mRMR | 0.746 (0.143) | 0.935 (0.010) | 0.928 (0.031) | 0.952 (0.009) | 0.712 (0.122) | 0.976 (0.004) | 0.968 (0.009) | 0.974 (0.015) |
| Pre-post treatment | WGCNA | - | 0.700 (0.018) | 0.696 (0.019) | 0.696 (0.019) | - | 0.649 (0.007) | 0.647 (0.005) | 0.645 (0.006) |
|  | ANOVA | - | 0.721 (0.058) | 0.679 (0.041) | 0.700 (0.063) | - | 0.733 (0.076) | 0.646 (0.052) | 0.680 (0.081) |
|  | mRMR | - | **0.779 (0.037)** | 0.675 (0.070) | 0.689 (0.034) | - | **0.786 (0.036)** | 0.660 (0.069) | 0.671 (0.043) |

**Bold values** indicate the best performance for each dataset.

### 3.2 Computational cost of GNNs

Table 2 documents the computational time for each model across different datasets. The GNN-based approaches required significantly more computational resources compared to traditional machine learning models. However, enhanced classification performance can justify this increased computational cost. Notably, our GNN implementations were run sequentially due to resource limitations; parallel processing could potentially lead to substantial improvements in computational efficiency.

**Table 2:** Time cost for different models across tasks.

| Model | Primary-recurrent | Normal-tumor | Pre-post treatment |
| --- | --- | --- | --- |
| Logistic | 0.5 s | 0.5 s | 0.4 s |
| Elastic net | 1.7 s | 3.1 s | 1.2 s |
| Random forest | 15.6 s | 2.6 s | 1.6 s |
| SVM | 6.8 s | 5.0 s | 1.9 s |
| MLP | 13.7 s | 7.4 s | 3.1 s |
| GCN-gene | 1~2 days | 2~4 hours | - |
| GCN-all-feature | 1~2 days | 2~4 hours | 2~3 hours |
| GAT-all-feature | 2~3 days | 3~6 hours | 3~6 hours |

GNN model performance could be further enhanced through two key optimizations: (1) implementation of a more comprehensive hyperparameter search strategy, and (2) adoption of more stringent thresholds for shared sample selection. These potential enhancements suggest that the current performance metrics of GNNs represent a conservative estimate of the methodology’s capabilities.

### 3.3 Graph-based features enhance performance

We utilized graph-based feature profiles to construct graph networks. The GNN predictions leveraging these networks demonstrated superior performance compared to all other feature types (Figure 4). In traditional machine learning models, graph-based features achieved perfect separation in the normal-tumor classification task and exceeded 90% accuracy and AUC in pre- and post-treatment sample classification. These features also delivered competitive performance for primary-recurrent classification.

**Figure 4:**
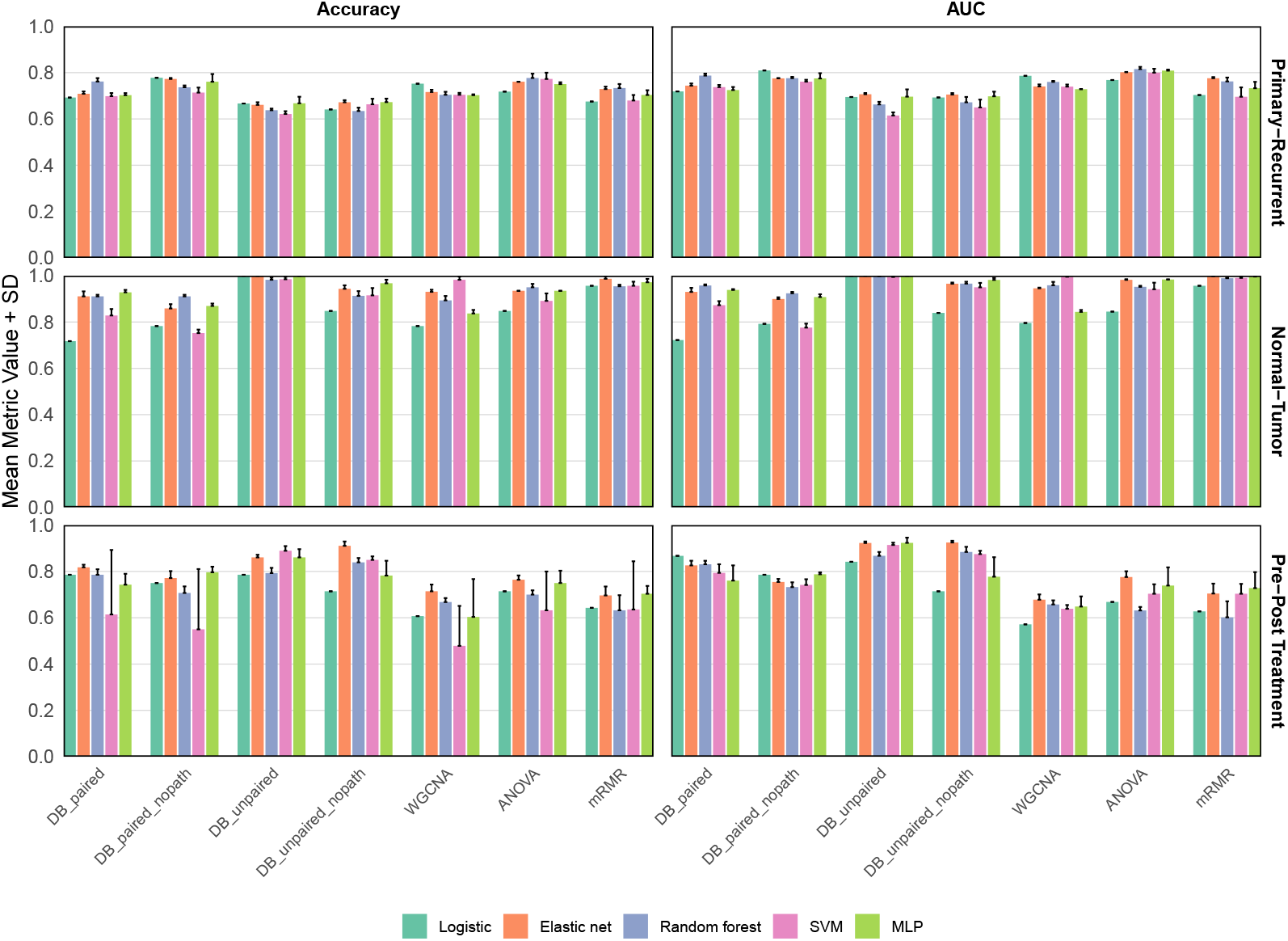
Performance of different features including graph-based features

The GNN models achieved an accuracy of 0.948 and an AUC of 0.977 in the primary-recurrent dataset, perfect separation in the normal-tumor dataset, and an accuracy of 0.896 with an AUC of 0.926 in the pre-post treatment dataset (Table 3). With our feature selection methodology, accuracy improved by 7.2%, 1.3%, and 16.9% in these respective tasks, while AUC increased by 2.4%, 0.2%, and 17.8%. The exceptional performance of DB_unpaired and DB_unpaired_nopath features in GNN models highlights the effectiveness of our approach in capturing complex biological relationships.

**Table 3:** Extended performance comparison with paired/unpaired datasets and feature selection methods.

| Dataset | Features | Accuracy (SD) |  |  |  | AUC (SD) |  |  |  |
| --- | --- | --- | --- | --- | --- | --- | --- | --- | --- |
|  |  | GCN-gene | GCN | GAT | GATv2 | GCN-gene | GCN | GAT | GATv2 |
| Primary-recurrent | DB_Paired | 0.816 (0.023) | 0.775 (0.047) | 0.733 (0.033) | 0.779 (0.014) | 0.882 (0.014) | 0.833 (0.041) | 0.785 (0.042) | 0.843 (0.019) |
|  | DB_paired_nopath | 0.821 (0.015) | 0.812 (0.037) | 0.733 (0.025) | 0.784 (0.052) | 0.896 (0.014) | 0.872 (0.035) | 0.773 (0.036) | 0.837 (0.063) |
|  | DB_unpaired | 0.938 (0.048) | 0.942 (0.014) | 0.915 (0.027) | 0.908 (0.016) | 0.977 (0.031) | **0.977 (0.015)** | 0.963 (0.015) | 0.958 (0.014) |
|  | DB_unpaired_nopath | **0.948 (0.016)** | 0.937 (0.023) | 0.910 (0.023) | 0.928 (0.019) | 0.971 (0.014) | 0.974 (0.024) | 0.961 (0.013) | 0.970 (0.012) |
|  | WGCNA | 0.717 (0.017) | 0.748 (0.058) | 0.698 (0.063) | 0.740 (0.008) | 0.780 (0.013) | 0.782 (0.085) | 0.731 (0.088) | 0.771 (0.011) |
|  | ANOVA | 0.838 (0.006) | 0.847 (0.034) | 0.855 (0.017) | 0.850 (0.020) | 0.901 (0.005) | 0.897 (0.032) | 0.901 (0.019) | 0.900 (0.024) |
|  | mRMR | 0.884 (0.010) | 0.820 (0.056) | 0.866 (0.021) | 0.862 (0.020) | 0.954 (0.004) | 0.892 (0.055) | 0.932 (0.018) | 0.924 (0.009) |
| Normal-tumor | DB_paired | 0.617 (0.104) | 0.911 (0.036) | 0.889 (0.047) | 0.899 (0.041) | 0.492 (0.264) | 0.944 (0.033) | 0.887 (0.031) | 0.919 (0.040) |
|  | DB_paired_nopath | 0.580 (0.060) | 0.902 (0.021) | 0.828 (0.033) | 0.885 (0.069) | 0.457 (0.168) | 0.940 (0.007) | 0.834 (0.034) | 0.903 (0.077) |
|  | DB_unpaired | 0.815 (0.033) | 0.980 (0.007) | **1.000 (0.000)** | 0.985 (0.034) | 0.832 (0.048) | 0.995 (0.004) | **1.000 (0.000)** | 0.993 (0.018) |
|  | DB_unpaired_nopath | 0.717 (0.093) | 0.948 (0.011) | 0.835 (0.122) | 0.889 (0.019) | 0.693 (0.121) | 0.960 (0.004) | 0.854 (0.132) | 0.917 (0.034) |
|  | WGCNA | 0.813 (0.027) | 0.963 (0.023) | 0.917 (0.030) | 0.922 (0.026) | 0.806 (0.014) | 0.991 (0.008) | 0.961 (0.020) | 0.967 (0.017) |
|  | ANOVA | 0.583 (0.073) | 0.943 (0.046) | 0.865 (0.132) | 0.913 (0.023) | 0.399 (0.191) | 0.978 (0.032) | 0.907 (0.144) | 0.950 (0.015) |
|  | mRMR | 0.746 (0.143) | 0.935 (0.010) | 0.928 (0.031) | 0.952 (0.009) | 0.712 (0.122) | 0.976 (0.004) | 0.968 (0.009) | 0.974 (0.015) |
| Pre-post | DB_paired | - | 0.793 (0.028) | 0.721 (0.041) | 0.711 (0.064) | - | 0.833 (0.030) | 0.740 (0.071) | 0.691 (0.064) |
|  | DB_paired_nopath | - | 0.793 (0.073) | 0.718 (0.052) | 0.696 (0.030) | - | 0.789 (0.080) | 0.698 (0.057) | 0.669 (0.050) |
|  | DB_unpaired | - | 0.846 (0.041) | 0.850 (0.015) | 0.843 (0.066) | - | 0.892 (0.034) | 0.886 (0.019) | 0.858 (0.092) |
|  | DB_unpaired_nopath | - | 0.796 (0.024) | 0.871 (0.063) | **0.896 (0.057)** | - | 0.839 (0.032) | 0.915 (0.053) | **0.926 (0.057)** |
|  | WGCNA | - | 0.700 (0.018) | 0.696 (0.019) | 0.696 (0.019) | - | 0.649 (0.007) | 0.647 (0.005) | 0.645 (0.006) |
|  | ANOVA | - | 0.721 (0.058) | 0.679 (0.041) | 0.700 (0.063) | - | 0.733 (0.076) | 0.646 (0.052) | 0.680 (0.081) |
|  | mRMR | - | 0.779 (0.037) | 0.675 (0.070) | 0.689 (0.034) | - | 0.786 (0.036) | 0.660 (0.069) | 0.671 (0.043) |

**Bold values** indicate the best performance for each dataset.

### 3.4 Biomarkers selected from unpaired approach

Our approach successfully identified key predictive markers from unpaired feature selection, many of which have literature support for their functional relevance (Supplementary table 1). In IDH-wt glioma, PGR, COL17A1, and RGS14 showed strong predictive value for recurrence, consistent with their known roles in tumor aggressiveness and therapy resistance (Cortese et al., 2021; Yan et al., 2022; Yang et al., 2024). For breast cancer normal-tumor differentiation, CXCL14, MSLN, LCN2, and MED1 emerged as significant predictors, supported by evidence of their differential expression and involvement in tumor progression (Yang et al., 2009; Gu et al., 2012; Tchou et al.; 2012; Zhang et al., 2013). In HER2-negative breast cancer treated with Bevacizumab, MMP2, COL5A2, COL6A1, DLK1, and MDK were identified as predictive of treatment response, aligning with their documented roles in angiogenesis and stromal interactions (Radenkovic et. al., 2014; Nueda et al., 2017; Li et. al., 2022; Gao et. al., 2023; Luo et al., 2025). These findings validate our biomarker discovery method and underscore the biological relevance of the identified markers, enhancing their potential translational value in precision oncology.

## 4 Discussion

The EGNF leverages the power of GNNs to advance biomarker identification through gene expression data, offering a sophisticated approach to modeling biological relationships. The predictive markers identified by EGNF across the three datasets, IDH-wt glioma, normal-tumor breast cancer, and HER2-negative breast cancer treated with Bevacizumab, exemplify the model’s ability to uncover biologically significant genes within complex expression networks.

In IDH-wt glioma, markers like PGR, COL17A1, and RGS14 reflect pathways tied to recurrence, such as progesterone signaling, extracellular matrix remodeling, and G-protein regulation, respectively, which are critical in aggressive glioma biology. Recurrence represents a major clinical challenge in glioma management, with IDH-wildtype tumors showing particularly aggressive behavior and high recurrence rates, often leading to treatment failure and poor patient outcomes (Weller et al., 2017; Molenaar et al., 2018). The identification of molecular signatures associated with recurrence could enable earlier intervention and personalized treatment strategies, potentially improving the dismal prognosis currently associated with these tumors.

For breast cancer normal-tumor differentiation, CXCL14, MSLN, LCN2, and MED1 highlight immune modulation, cell surface alterations, inflammation, and estrogen receptor coactivation, processes central to tumorigenesis and supported by differential expression evidence. In the Bevacizumab-treated HER2-negative breast cancer cohort, markers MMP2, COL5A2, COL6A1, DLK1, and MDK illustrate angiogenesis regulation, stromal architecture, and growth factor signaling pathways. Treatment resistance in breast cancer, particularly to targeted therapies like Bevacizumab, remains a significant obstacle in clinical management, with most patients eventually developing resistance mechanisms that lead to disease progression (Bergers & Hanahan, 2008). MMP2 and collagen genes influence extracellular matrix dynamics affecting drug delivery, while MDK provides alternative angiogenic signaling potentially contributing to treatment resistance. These markers represent interconnected biological processes that determine Bevacizumab response, offering insights for patient stratification in anti-angiogenic therapy and potentially enabling clinicians to identify patients who might benefit from alternative or combination approaches before resistance develops.

Importantly, EGNF demonstrates two significant advantages that extend beyond our specific test cases. First, it offers a powerful approach for biomarker identification, extracting genes with genuine biological relevance rather than statistical artifacts. This capability stems from the method’s integration of network topology with expression data, enabling it to capture complex gene interactions that traditional feature selection methods might miss. Second, EGNF provides superior predictive performance across various classification tasks, as evidenced by our comparative analyses. This dual strength, identifying meaningful biomarkers while enhancing prediction accuracy, makes EGNF particularly valuable in clinical applications where both mechanistic insights and reliable patient stratification are crucial. Moreover, these capabilities suggest that EGNF could be effectively applied to any sample classification task involving gene expression data, regardless of the disease context or research.

Recent developments in the field provide valuable insights into how GNN-based methods, like EGNF, can be contextualized and potentially enhanced. One notable trend is the application of GNNs to multi-modal biological data for improved disease classification and biomarker discovery. Researchers have explored hybrid GNN models that integrate gene expression profiles with protein-protein interaction networks to identify cancer-specific biomarkers, achieving high predictive accuracy by capturing cross-modal dependencies (Rhee et al., 2018). This suggests that EGNF could benefit from incorporating additional data types, such as epigenetic or proteomic information, to enrich its network representations. The ability to model such complex interactions is a key advantage of GNNs, enabling the detection of subtle patterns that traditional methods might overlook.

Another area of progress is the use of GNNs for rare disease research, where data scarcity poses a significant challenge. Studies have shown that GNN-based frameworks, combined with transfer learning, can effectively identify biomarkers using limited gene expression datasets (Dutil et al., 2018; Theodoris et al., 2023). This is particularly relevant for EGNF, as its performance in scenarios with small sample sizes, such as distinguishing nuanced disease states, could be bolstered by adopting similar techniques. By pre-training on larger, related datasets and fine-tuning on specific tasks, EGNF might overcome limitations associated with data availability, enhancing its applicability in precision medicine.

Interpretability remains a critical focus in GNN research, aligning with EGNF’s emphasis on biologically meaningful feature selection. Advances have introduced GNN models with explainability layers that highlight key gene sub-networks driving predictions in cancer prognosis (Ying et al., 2019; Chereda et al., 2021). This approach not only improves classification but also provides clinicians with actionable insights into disease mechanisms. Integrating such interpretability mechanisms into EGNF could strengthen its capacity to identify significant gene modules, making it more valuable for translational research where biological relevance is paramount. Scalability is another consideration, as GNNs often require substantial computational resources. Research has demonstrated that optimization techniques, such as pruning redundant graph connections, can reduce computational overhead without sacrificing accuracy in gene expression tasks (Chen et al., 2020). For EGNF, adopting such strategies could make it more feasible for large-scale genomic studies, broadening its practical utility.

In conclusion, EGNF stands to gain from these emerging trends in GNN research, including multi-modal integration, transfer learning for small datasets, enhanced interpretability, and computational optimization. These advancements highlight the potential for EGNF to evolve into an even more robust tool for biomarker identification and accurate prediction, bridging the gap between computational power and clinical impact in addressing critical challenges like glioma recurrence and breast cancer treatment resistance.

## Acknowledgments

This work was supported by the MD Anderson Moonshot Program. We also want to thank the kind comments and suggestions from Dr. Xi Luo, Dr. Vahed Maroufy and Dr. Goo Jun at UT Health Houston.

## Author contributions

Y.L., K.K., and J.H. conceived and designed the study. Y.L. performed the computational analysis and wrote the manuscript. K.K., and J.H. edited the manuscript. All the authors reviewed and approved the final manuscript.

## Data Availability

The gene expression datasets used in this study are publicly available from the GLASS Consortium (https://www.synapse.org/Synapse:syn17038081/wiki/585622), TCGA (https://www.cancer.gov/ccg/research/genome-sequencing/tcga), and the GEO under accession number GSE87455 (https://www.ncbi.nlm.nih.gov/geo/query/acc.cgi?acc=GSE87455).

## Code availability

The source code and tutorial of this work can be found on GitHub https://github.com/yliu38/EGNF.

## Notes

### Competing Interest Statement

The authors have declared no competing interest.

### Summary of Updates

replace old references with new ones.

## References

[1] Bergers, G., & Hanahan, D. (2008). Modes of resistance to anti-angiogenic therapy. Nature Reviews Cancer, 8(8), 592–603. 10.1038/nrc2442

[2] Bergstra, J., & Bengio, Y. (2012). Random search for hyper-parameter optimization. Journal of Machine Learning Research, 13(1), 281–305. https://jmlr.org/papers/v13/bergstra12a.html

[3] Breiman, L. (2001). Random forests. Machine Learning, 45(1), 5–32. 10.1023/A:010933404324

[4] Brody, S., Alon, U., & Yahav, E. (2022). How attentive are graph attention networks? arXiv. 10.48550/arXiv.2105.14491

[5] Chen, J., Ma, T., & Xiao, C. (2020). FastGCN: Fast learning with graph convolutional networks via importance sampling. International Conference on Learning Representations. https://arxiv.org/abs/1801.10247

[6] Chereda, H., Bleckmann, A., Menck, K., Perera-Bel, J., Stegmaier, P., Auer Anonymous, F., Kramer, F., Leha, A., & Beißbarth, T. (2021). Explaining decisions of graph convolutional neural networks: Patient-specific molecular subnetworks responsible for metastasis prediction in breast cancer. Genome Medicine, 13(1), 42. 10.1186/s13073-021-00845-7

[7] Cortes, C., & Vapnik, V. (1995). Support-vector networks. Machine Learning, 20(3), 273–297. 10.1007/BF00994018

[8] Cortese, A. C., Smart, C., & Brown, S. D. (2021). Intracellular progesterone receptor and cSrc protein working together to promote migration and invasion of human glioblastoma cells. Frontiers in Endocrinology, 12, 640298. 10.3389/fendo.2021.640298

[9] Dutil, F., Cohen, J. P., Weiss, M., Derevyanko, G., & Bengio, Y. (2018). Towards gene expression convolutions using gene interaction graphs. International Conference on Machine Learning, Workshop on Computational Biology. https://arxiv.org/abs/1806.06975

[10] Gao, S., Wang, Y., Xu, Y., & Liu, S. (2023). An angiogenesis-related lncRNA signature is associated with prognosis and tumor immune microenvironment in breast cancer. Journal of Personalized Medicine, 13(3), 513. 10.3390/jpm13030513

[11] Gu, X. L., Ou, Z. L., Lin, F. J., Yang, X. L., Luo, J. M., Shen, Z. Z., & Shao, Z. M. (2012). Expression of CXCL14 and its anticancer role in breast cancer. Breast Cancer Research and Treatment, 135(3), 725–735. 10.1007/s10549-012-2206-2

[12] Hosmer, D. W., Lemeshow, S., & Sturdivant, R. X. (2013). Applied logistic regression (3rd ed.). Wiley. 10.1002/9781118548387

[13] Kipf, T. N., & Welling, M. (2017). Semi-supervised classification with graph convolutional networks. arXiv. 10.48550/arXiv.1609.02907

[14] Kuhn, M. (2008). Building predictive models in R using the caret package. Journal of Statistical Software, 28(5), 1–26. 10.18637/jss.v028.i05

[15] Langfelder, P., & Horvath, S. (2008). WGCNA: An R package for weighted correlation network analysis. BMC Bioinformatics, 9, 559. 10.1186/1471-2105-9-559

[16] Li, X., Li, Z., Gu, S., & Zhao, X. (2022). A pan-cancer analysis of collagen VI family on prognosis, tumor microenvironment, and its potential therapeutic effect. BMC Bioinformatics, 23(1), 390. 10.1186/s12859-022-04951-0

[17] Luo, L., Yang, P., Mastoraki, S., Rao, X., Wang, Y., Kettner, N. M., Raghavendra, A. S., Tripathy, D., Damodaran, S., Hunt, K. K., Wang, J., Li, Z., & Keyomarsi, K. (2025). Single-cell RNA sequencing identifies molecular biomarkers predicting late progression to CDK4/6 inhibition in patients with HR+/HER2-metastatic breast cancer. Molecular Cancer, 24(1), 48. 10.1186/s12943-025-02226-9

[18] MacQueen, J. B. (1967). Some methods for classification and analysis of multivariate observations. In L.M. Le Cam & J. Neyman (Eds.), Proceedings of the fifth Berkeley symposium on mathematical statistics and probability (Vol. 1, pp. 281–297). University of California Press. https://projecteuclid.org/euclid.bsmsp/1200512992

[19] Molenaar, R. J., Maciejewski, J. P., Wilmink, J. W., & van Noorden, C. J. F. (2018). Wild-type and mutated IDH1/2 enzymes and therapy responses. Oncogene, 37(15), 1949–1960. 10.1038/s41388-017-0077-z

[20] Mostafa, S. M., & Amano, H. (2019). Effect of clustering data in improving machine learning model accuracy. Journal of Theoretical and Applied Information Technology, 97(21), 2973–2981. https://www.jatit.org/volumes/Vol97No21/7Vol97No21.pdf

[21] Nueda, M. L., Naranjo, A. I., Baladrón, V., & Laborda, J. (2017). Different expression levels of DLK1 inversely modulate the oncogenic potential of human MDA-MB-231 breast cancer cells through inhibition of NOTCH1 signaling. The FASEB Journal, 31(8), 3484–3496. 10.1096/fj.201601341RRR

[22] Radenkovic, S., Konjevic, G., Jurisic, V., Karadzic, K., Nikitovic, M., & Gopcevic, K. (2014). Values of MMP-2 and MMP-9 in tumor tissue of basal-like breast cancer patients. Cell Biochemistry and Biophysics, 68(1), 143–152. 10.1007/s12013-013-9701-x

[23] Ramirez, R., Chiu, Y. C., Hererra, A., Mostavi, M., Ramirez, J., Chen, Y., Huang, Y., & Jin, Y. F. (2020). Classification of cancer types using graph convolutional neural networks. Frontiers in Physics, 8, 203. 10.3389/fphy.2020.00203

[24] Rhee, S., Seo, S., & Kim, S. (2018). Hybrid approach of relation network and localized graph convolutional filtering for breast cancer subtype classification. Proceedings of the Twenty-Seventh International Joint Conference on Artificial Intelligence, 3527–3534. 10.24963/ijcai.2018/490

[25] Rumelhart, D. E., Hinton, G. E., & Williams, R. J. (1986). Learning representations by back-propagating errors. Nature, 323(6088), 533–536. 10.1038/323533a0

[26] Snoek, J., Larochelle, H., & Adams, R. P. (2012). Practical Bayesian optimization of machine learning algorithms. Advances in Neural Information Processing Systems, 25, 2951–2959. 10.48550/arXiv.1206.2944

[27] Tchou, J., Wang, L. C., Selven, B., Zhang, H., Conejo-Garcia, J., Borghaei, H., Kalos, M., Vondeheide, R. H., Albelda, S. M., June, C. H., & Zhang, L. (2012). Mesothelin, a novel immunotherapy target for triple negative breast cancer. Breast Cancer Research and Treatment, 133(2), 799–804. 10.1007/s10549-012-2018-4

[28] Theodoris, C. V., Xiao, L., Chopra, A., Chaffin, M. D., Al Sayed, Z. R., Hill, M. C., Mantineo, H., Brydon, E. M., Zeng, Z., Liu, X. S., & Ellinor, P. T. (2023). Transfer learning enables predictions in network biology. Nature, 618(7965), 616–624. 10.1038/s41586-023-06139-9

[29] Veličković, P., Cucurull, G., Casanova, A., Romero, A., Liò, P., & Bengio, Y. (2018). Graph attention networks. arXiv. 10.48550/arXiv.1710.10903

[30] Wang, T., Shao, W., Huang, Z., Tang, H., Zhang, J., Ding, Z., & Huang, K. (2021). MOGONET integrates multi-omics data using graph convolutional networks allowing patient classification and biomarker identification. Nature Communications, 12(1), 3445. 10.1038/s41467-021-23774-w

[31] Weller, M., van den Bent, M., Tonn, J. C., Stupp, R., Preusser, M., Cohen-Jonathan-Moyal, E., Henriksson, R., Le Rhun, E., Balana, C., Chinot, O., Bendszus, M., Reijneveld, J. C., Dhermain, F., French, P., Marosi, C., Watts, C., Oberg, I., Pilkington, G., Baumert, B. G., … Wick, W. (2017). European Association for Neuro-Oncology (EANO) guideline on the diagnosis and treatment of adult astrocytic and oligodendroglial gliomas. The Lancet Oncology, 18(6), e315–e329. 10.1016/S1470-2045(17)30194-8

[32] Yan, Y., Li, X., Zhang, Y., Dinnyés, A., & Zhang, Y. (2022). Integrated analysis of ECT2 and COL17A1 as potential biomarkers of glioblastoma multiforme. BioMed Research International, 2022, Article 9453549. 10.1155/2022/9453549

[33] Yang, J., Bielenberg, D. R., Rodig, S. J., Doiron, R., Clifton, M. C., Kung, A. L., Himanen, J. P., & Moses, M. A. (2009). Lipocalin 2 promotes breast cancer progression. Proceedings of the National Academy of Sciences, 106(10), 3913–3918. 10.1073/pnas.0810617106

[34] Yang, S., Wang, X., Huan, R., Deng, M., Kong, Z., Xiong, Y., Luo, T., Jin, Z., Liu, J., Chu, L., Han, G., Zhang, J., & Tan, Y. (2024). Machine learning unveils immune-related signature in multicenter glioma studies. iScience, 27(4), 109317. 10.1016/j.isci.2024.109317

[35] Ying, Z., Bourgeois, D., You, J., Zitnik, M., & Leskovec, J. (2019). GNNExplainer: Generating explanations for graph neural networks. Advances in Neural Information Processing Systems, 32, 9240–9251. https://proceedings.neurips.cc/paper/2019/file/d80b7040b773199015de6d3b4293c8ff-Paper.pdf

[36] Zhang, L., Cui, J., Leonard, M., Nephew, K., Li, Y., & Zhang, X. (2013). Silencing MED1 sensitizes breast cancer cells to pure anti-estrogen fulvestrant in vitro and in vivo. PLoS ONE, 8(7), e70641. 10.1371/journal.pone.0070641

[37] Zou, H., & Hastie, T. (2005). Regularization and variable selection via the elastic net. Journal of the Royal Statistical Society: Series B (Statistical Methodology), 67(2), 301–320. 10.1111/j.1467-9868.2005.00503.x

